# Deep Learning-Based Structure Modeling of the *Treponema pallidum* Proteome: Insights into Pathogenesis and Syphilis Vaccine Development

**DOI:** 10.64898/2026.05.05.717303

**Authors:** Simon Houston, Steven Marshall, Austin Miller, Aleksander Palkowski, Javier A. Alfaro, Caroline E. Cameron

## Abstract

*Treponema pallidum* ssp. *pallidum*, the causative agent of syphilis, has a small proteome and encompasses numerous strains. Knowledge gaps remain in understanding the molecular mechanisms of pathogenesis of this bacterium, as well as the structure and function of the full complement of proteins encoded by *T. pallidum*. Here, an AI-based structure-to-function modeling workflow was used to investigate the complement of proteins encoded by *T. pallidum*. High-confidence structure models were generated for 976 *T. pallidum* proteins, covering 99% of the proteome. Analysis of the generated models using the protein structure comparison server DALI enabled high-confidence, structure-based functional annotation of 877 *T. pallidum* proteins, including 240 of the 323 proteins of unknown function encoded by this pathogen. Additionally, 63 putative pathogenesis related proteins (PPRPs) and seven treponemal proteins with previously uncharacterized similarity to outer membrane proteins (OMPs) from Gram-negative bacteria were identified. A workflow for B cell epitope (BCE) prediction identified 1133 surface-exposed, host-facing potential epitopes in known and predicted *T. pallidum* OMPs, of which 92 were prioritized based on bioinformatic analyses, biophysical properties, amino acid sequence conservation, and previous protein expression data. This work provides insight into *T. pallidum* pathogenesis through structure modeling-based functional annotation, including characterization of proteins of unknown function. This study also informs syphilis vaccine design by identifying new potential *T. pallidum* OMPs, as well as host-facing regions of *T. pallidum* OMPs that have conserved amino acid sequences in globally circulating strains.

**Statement of importance/impact:** This study presents the first AI-based global structure modeling-to-function analysis of the proteome of *Treponema pallidum*, the bacterium that causes syphilis. Structure-based functional predictions of previously uncharacterized proteins, including proteins potentially involved in virulence, provide novel insight into mechanisms of pathogenesis. The work also informs syphilis vaccine development by the identification and structural characterization of new candidate vaccine proteins in globally circulating strains of *T. pallidum*.

## Introduction

*Treponema pallidum* ssp. *pallidum* (hereafter *T. pallidum*) is a bacterium that causes the chronic and multistage sexually transmitted infection syphilis and the vertically transmitted disease congenital syphilis. The continued increase in syphilis rates worldwide indicates that current public health strategies are insufficient for reducing disease incidence, highlighting the need for development of an effective syphilis vaccine (Aho et al. 2022; Arrieta and Singh 2019; Cameron and Lukehart 2014; Chen et al. 2023; Gilmour and Walls 2023; Korenromp et al. 2019; Spiteri et al. 2019; Tsuboi et al. 2021; Waugh and Cameron 2024).

All known strains of *T. pallidum* can be classified into two main phylogenetic clades, each named after their respective reference strains: the Nichols clade and the SS14 clade (Arora et al. 2016; Mikalova et al. 2010; Nechvatal et al. 2014; Petrosova et al. 2013; Smajs et al. 2012). SS14 strains exhibit a higher frequency of macrolide resistance (Stamm and Bergen 2000; Stamm et al. 1988), and according to current clinical sampling data are more prevalent worldwide (Arora et al. 2016; Beale et al. 2021; Lieberman et al. 2021; Sena et al. 2024). The *T. pallidum* genome consists of a 1.14 Mb single circular chromosome with approximately 1000 predicted protein-coding genes (Fraser et al. 1998). All *T. pallidum* genomes sequenced to date are highly conserved, with approximately 99.9% identity between whole genomes from different strains (Fraser et al. 1998; Matejkova et al. 2008; Petrosova et al. 2012; Smajs et al. 2012).

Despite this conservation, inter-strain genetic variation often results in amino acid sequence differences within predicted surface-exposed, host-facing regions of known and potential outer membrane proteins (OMPs) (Arora et al. 2016; Bettin et al. 2025; Houston et al. 2024b; Lieberman et al. 2021; Mikalova et al. 2010; Nechvatal et al. 2014; Petrosova et al. 2013; Pospisilova et al. 2025; Sena et al. 2024; Smajs et al. 2012). *T. pallidum* OMPs are the focus of current syphilis vaccine development strategies (Hawley et al. 2021; Izard et al. 2009; Liu et al. 2024; Liu et al. 2010; Radolf and Kumar 2017).

*Treponema pallidum* is a phylogenetically distinct bacterium, with approximately 30% of its predicted protein-coding genes lacking known orthologs and/or assigned functions (Fraser et al. 1998; Petrosova et al. 2013). Additionally, the genome contains few homologs of established virulence factors found in more conventional bacterial pathogens (Fraser et al. 1998; Petrosova et al. 2013). Proteomics studies have detected the expression of 95% of the *T. pallidum* proteome, including approximately 90% of the proteins of unknown function. This protein category encompasses most of the proteins annotated as “hypothetical”, as well as the majority of known and potential OMPs and predicted pathogenesis-related proteins (PPRPs) (Houston et al. 2023; Houston et al. 2024a; Houston et al. 2024b; McGill et al. 2010; Osbak et al. 2016). Although these proteins may contribute to the unique biology and pathogenesis of *T. pallidum* (Houston et al. 2018), knowledge of their functions remains limited.

Tertiary structure, which is more conserved than primary structure and directly influences how a protein carries out its specific biological role, is key to determining protein function (Baker and Sali 2001; Illergard et al. 2009). Therefore, structural biology approaches can be used to provide insights into protein function, which is especially relevant for proteins from experimentally intractable pathogens or pathogens expressing proteins that lack significant amino acid sequence orthology. However, only 20 structures from different *T. pallidum* proteins have been solved and deposited in the Protein Data Bank (PDB) (Berman et al. 2000; Brautigam et al. 2015; 2016; Brautigam et al. 2017; Brautigam et al. 2012; Brautigam et al. 2023; Deka et al. 2013a; Deka et al. 2012; Deka et al. 2013b; Deka et al. 2007; Deka et al. 2006; Deka et al. 2020; Deka et al. 2002; Deka et al. 2004; Lee et al. 1999; Luthra et al. 2011; Machius et al. 2007; Parker et al. 2016; Ramaswamy et al. 2019; Santos-Silva et al. 2006; Thumiger et al. 2006), representing just 2% of the *T. pallidum* proteome. To date, the structure of only one *T. pallidum* protein with experimental evidence supporting an outer membrane location has been solved: the vascular adhesin and syphilis vaccine candidate, TPANIC_0751 (Lithgow et al. 2017; Lithgow et al. 2021; Lukehart et al. 2022; Parker et al. 2016). In the absence of amino acid sequence orthology, functional annotations, and solved protein structures, the use of computational modeling enables high-throughput analyses of protein structure-function relationships and the prediction of protein functions.

Deep learning-based modeling programs, including AlphaFold (Abramson et al. 2024; Varadi et al. 2022) and RoseTTAFold (Baek et al. 2021), have transformed the field of protein structure prediction by providing higher modeling accuracy, including for proteins with low or no sequence homology (Baek et al. 2021; Jumper et al. 2021; Jumper and Hassabis 2022; Senior et al. 2020; Wang et al. 2024). These artificial intelligence (AI) modeling programs are not reliant on the availability of homologous protein structures. Instead, the amino acid sequence of the query protein and multiple sequence alignments (MSAs) are used to infer evolutionary information between amino acid residue pairs, enabling prediction of three-dimensional atomic-level structures with near-experimental accuracy (Baek et al. 2021; Baek et al. 2024; Jumper et al. 2021; Jumper and Hassabis 2022; Senior et al. 2020).

This study advances understanding of *T. pallidum* pathogenesis by assigning AI structure-based functional annotations to *T. pallidum* proteins of unknown function and identifying treponemal proteins with pathogenesis-related predicted functions. Further, the characterization of seven novel putative OMPs, and the prediction of B cell epitopes from the complement of OMPs, informs syphilis vaccine design.

## Results

### Proteome-wide distribution of *T. pallidum* protein structure modeling confidence scores

Using RoseTTAFold, 958 of the 990 *T. pallidum* proteins were modeled with confidence scores ≥ 0.5 (50%), with an average confidence score of 0.77 (**Figures 1 and 2 and Table S1)**. Re-analysis of the 32 low-confidence or unmodeled proteins using I-TASSER (Yang et al. 2015; Yang and Zhang 2015a; 2015b) or the AlphaFold database (Varadi et al. 2022; Varadi et al. 2024) yielded 18 additional high-confidence structure models (**Tables S1 and S3**). Overall, high-confidence structure models were obtained for 976 of the 990 *T. pallidum* proteins (**Table S1**).

**Figure 1:**
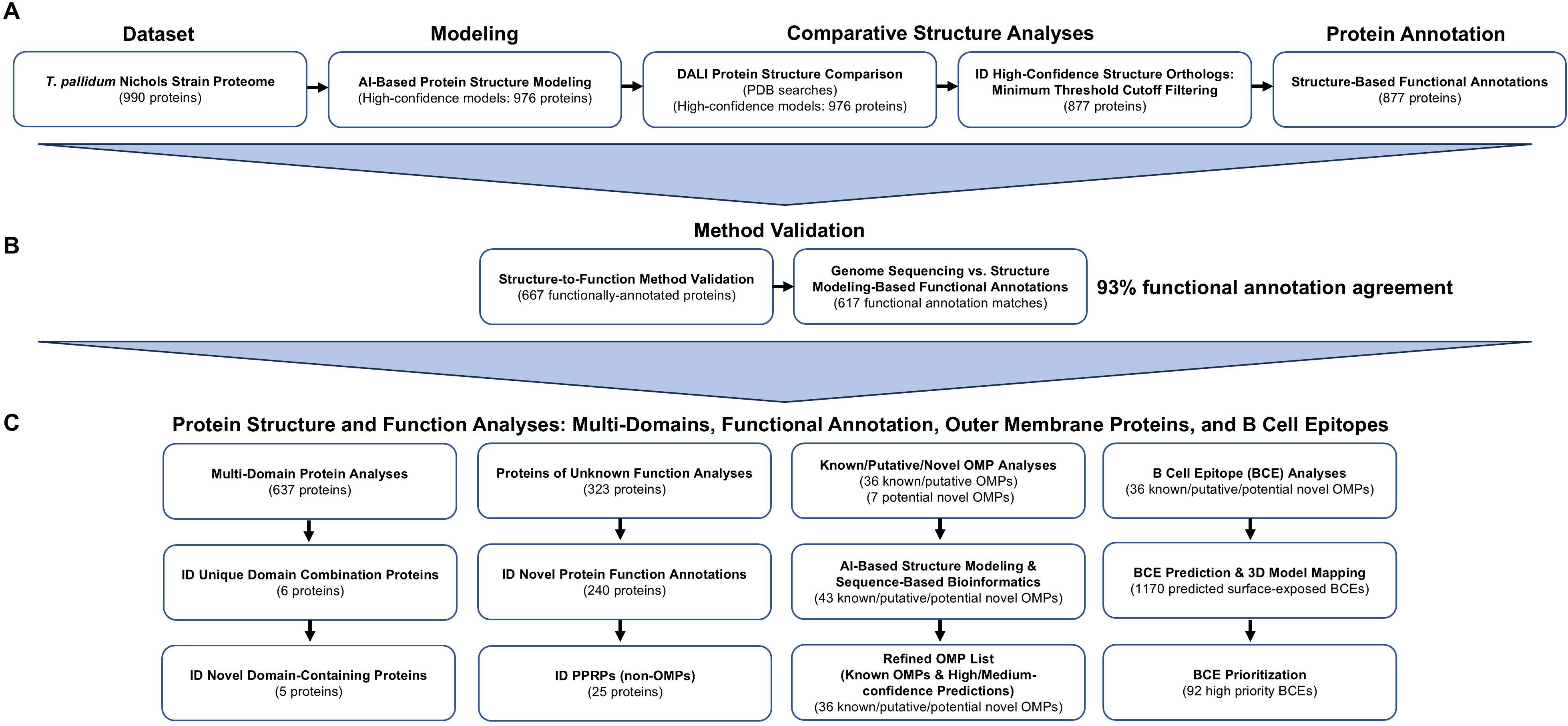
Structure modeling-to-function workflow. (A) 990 *T. pallidum* (Nichols strain) proteins were submitted to RoseTTAFold (RF). Models with confidence scores of at least 0.5 were analyzed using DALI for the identification of high-confidence structural orthologs in the PDB and the assignment of predicted protein functions. Of the 976 high-confidence structure models, 18 were generated using I-TASSER or sourced from the AlphaFold database due to RF limitations/low confidence scores. (B) Method validation was performed by comparing the functional annotations of 667 *T. pallidum* proteins with the corresponding structure-based functions predicted in the present study. (C) An array of protein structure and functional analyses were performed to gain insight into *T. pallidum* pathogenesis and to inform syphilis vaccine design by investigating multi-domain proteins, protein function, known and novel OMPs, and the identification and prioritization of BCEs.

**Figure 2.**
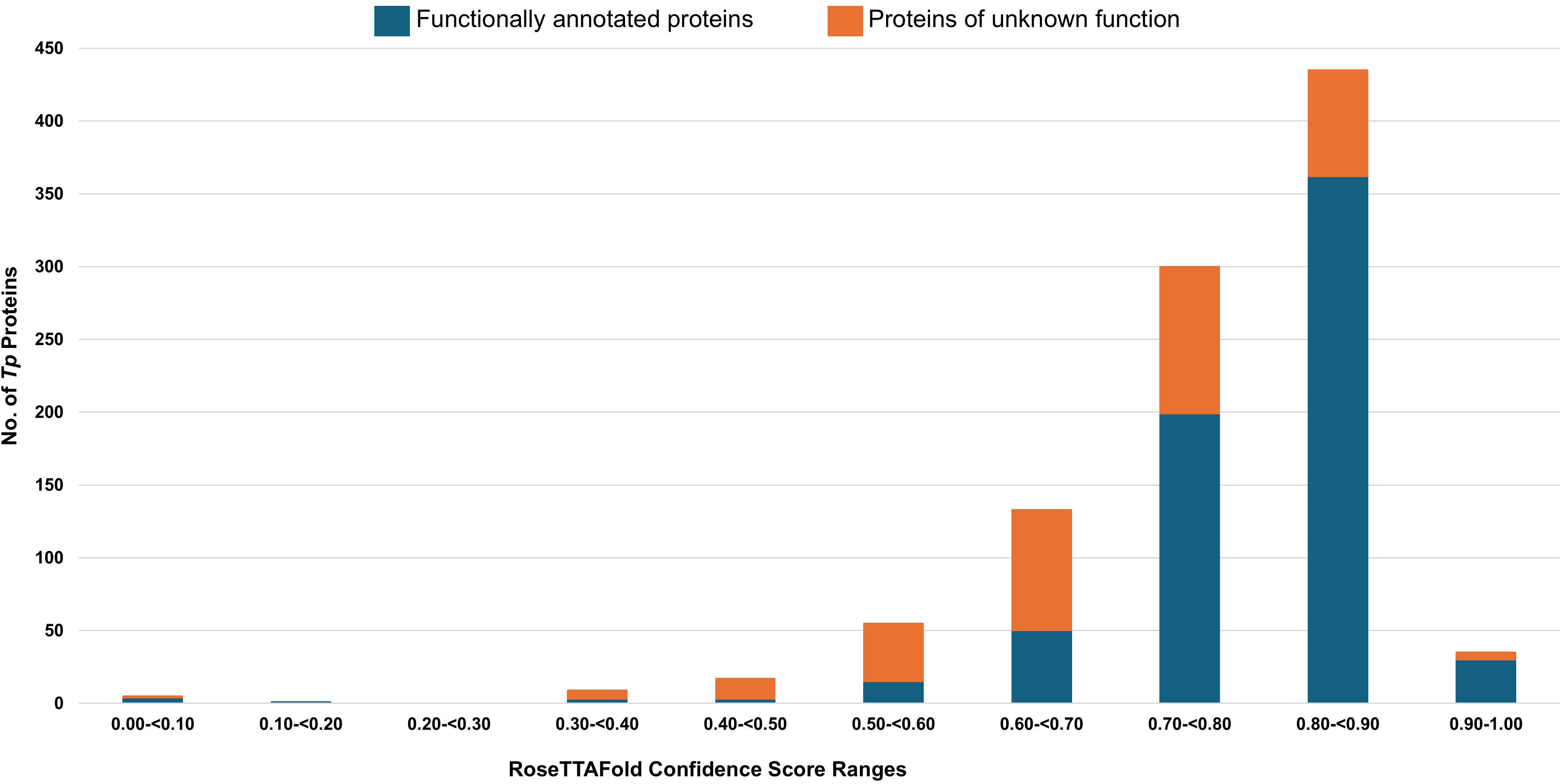
Distribution of RoseTTAFold confidence scores from structure modeling of the *T. pallidum* proteome. Bar graph depicting the distribution of *T. pallidum* proteins based on their RoseTTAFold confidence scores. A minimum confidence threshold score of 0.5 was used for further analyses, including the identification of high-confidence *T. pallidum* protein structure model orthologs. The number of functionally annotated proteins and proteins of unknown function are shown in blue and orange, respectively. The five proteins in the 0.00-<0.10 confidence range could not be modeled in RoseTTAFold due to protein size constraints.

### Identification of high-confidence *T. pallidum* protein structure model orthologs

Following filtering of the DALI data using stringent confidence thresholds, high-confidence structural orthologs were identified for 877 *T. pallidum* proteins, corresponding to approximately 89% of the proteome (**Figure 3A and Table S2)**. In general, the top-ranking, high-confidence DALI structure ortholog matches for functionally annotated *T. pallidum* proteins had higher Z-scores and alignment coverages compared to those of proteins of unknown function (**Figure 3B and Tables S2 and S4**). Overall, high Z-scores and alignment coverages were obtained for most structural orthologs, supporting high-confidence structure-based functional inference for most *T. pallidum* proteins.

**Figure 3.**
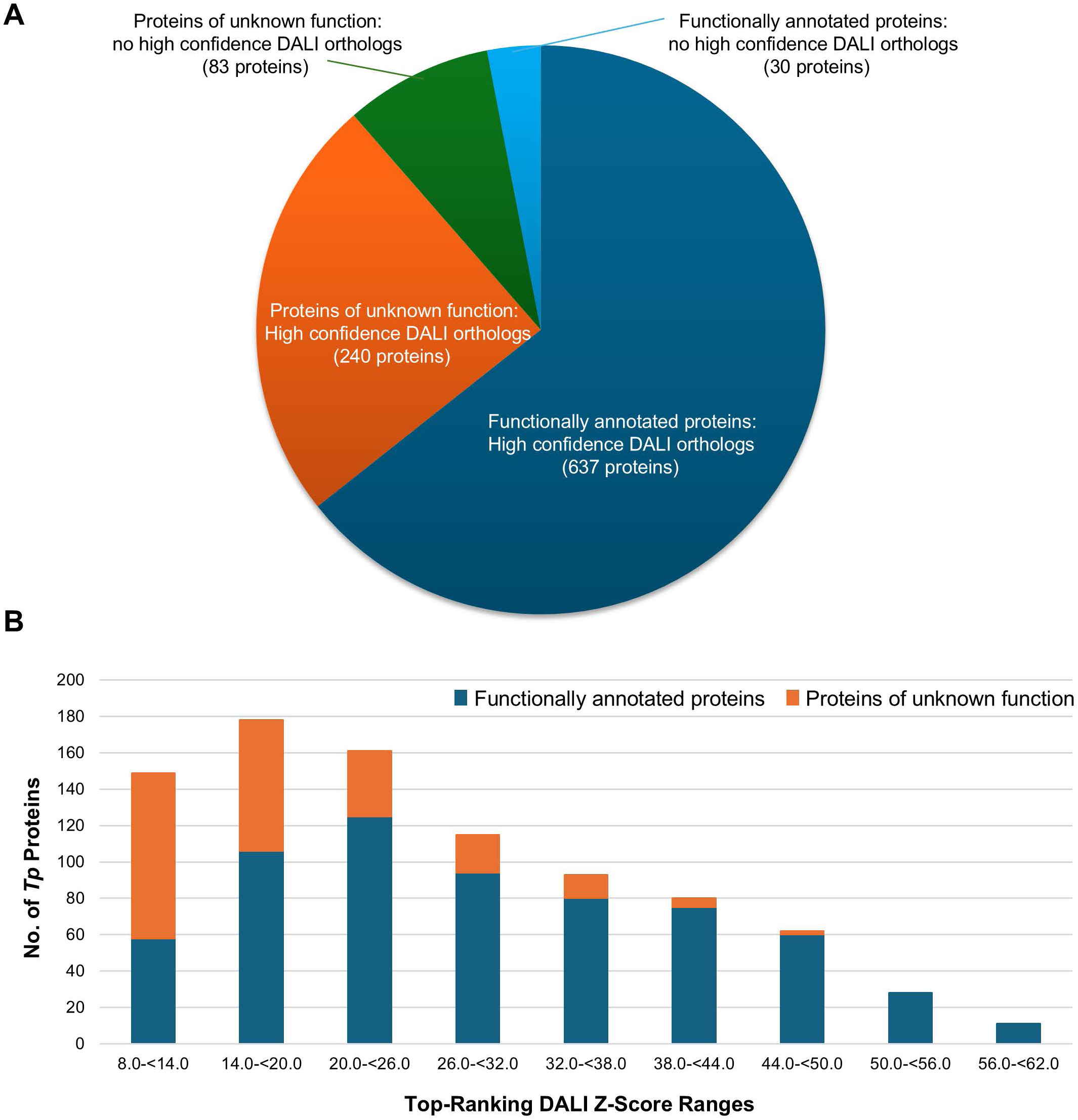
High-confidence *T. pallidum* protein structure model orthologs. (A) Pie chart showing the number of treponemal proteins that met or failed the minimum threshold scores for the identification of high-confidence DALI structural orthologs. (B) Bar graph illustrating the distribution of the top-ranking DALI Z-scores for each of the 877 treponemal proteins that were identified with high-confidence DALI structural orthologs. The number of functionally annotated proteins and proteins of unknown function are shown in blue and orange, respectively.

### Method validation: *T. pallidum* protein structure modeling-to-function pipeline

In the present AI-based modeling study, we compared genome sequence- and structure modeling-based functional annotations to validate our structure-to-function methodology. Out of a total of 667 *T. pallidum* proteins with genome sequencing-based functional annotations, 553 proteins were assigned the same function via our structure modeling pipeline (**Figure 4A and Table S5_part 1**). Additionally, 72 of these 553 protein structure modeling-based annotations provided more functional specificity when compared to their corresponding genome sequencing annotations (**Table S5_part 2**). A further 64 treponemal proteins were assigned structure-based functions that were similar to their genomic annotations (**Figure 4A and Table S5_part 3**). Only 10 proteins were assigned high-confidence functions in the present study that differed from their genome-based functional annotations (**Figure 4A and Table S5_part 4**). The remaining 40 proteins could not be assigned specific structure modeling-based functions and were omitted from the analysis (**Figure 4A and Table S5_parts 5 and 6**). COG analysis showed that most *T. pallidum* proteins with matching genome- and structure-based functional annotations were categorized as proteins with functions related to “Translation, ribosomal structure and biogenesis”, “Cell wall/membrane/envelope biogenesis”, and “Replication, recombination and repair”, whereas the COG classification for several proteins with differing functional annotations was “Function unknown” (**Figure 4B and Table S6**). Together, these findings demonstrate a strong correlation between genome and AI-based structure modeling functional annotations, which supports the validation of our structure modeling-to-function workflow.

**Figure 4.**
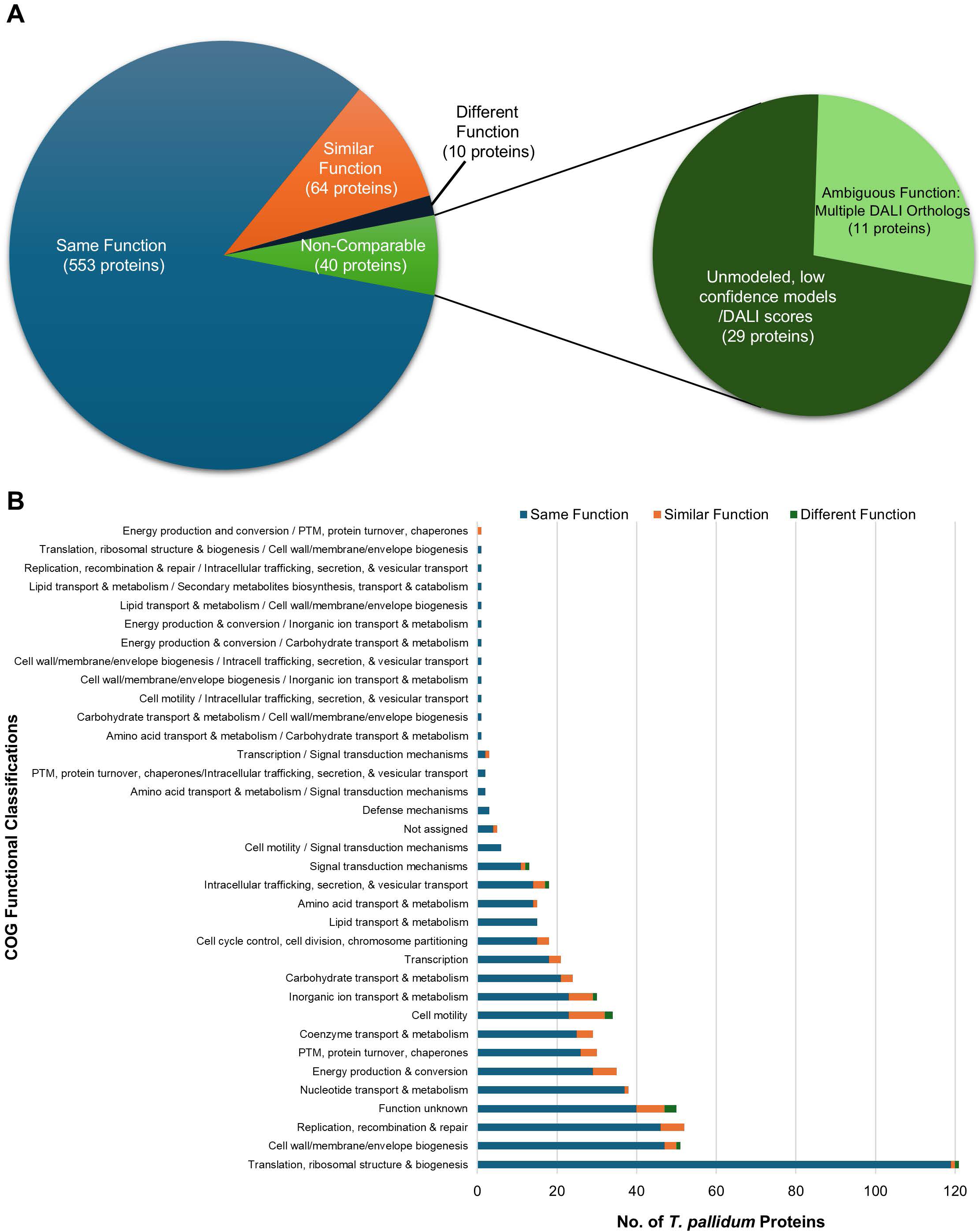
Method validation: NCBI proteome vs. AI-based modeling functional annotation comparisons. (A) Pie chart (left) depicting the number of treponemal proteins with structure modeling-based functional annotations that are the same, similar, or different compared to the functions assigned by the NCBI proteome annotation pipeline. The distribution of *T. pallidum* proteins with functions that could not be compared between the two methods is shown in the inset pie chart (right). (B) Bar chart showing the COG-based functional classification distribution of *T. pallidum* modeled proteins predicted to have the same, similar, or different functions compared to the NCBI proteome functional annotations. “Not assigned” COG functional classification – *T. pallidum* proteins for which eggNOG-mapper was unable to assign COG-based functional classifications.

### Identification of functionally annotated *T. pallidum* proteins with unique domain combinations and domains of unknown function

Structure models and DALI ortholog data for all 637 functionally annotated *T. pallidum* proteins with high-confidence structural orthologs were analyzed to identify proteins with unique domain combinations and domains of unknown function. Six proteins were identified with domain combinations not previously observed within the same protein in other species (**Figures 1C and 5A and Tables S7 and S8_part 1**). Five multi-domain *T. pallidum* proteins were also found to contain at least one domain that lacked significant structural orthology to any other protein domains identified in the DALI PDB searches (**Figure 1C and 5B and Tables S7 and S8_part 2**).

**Figure 5.**
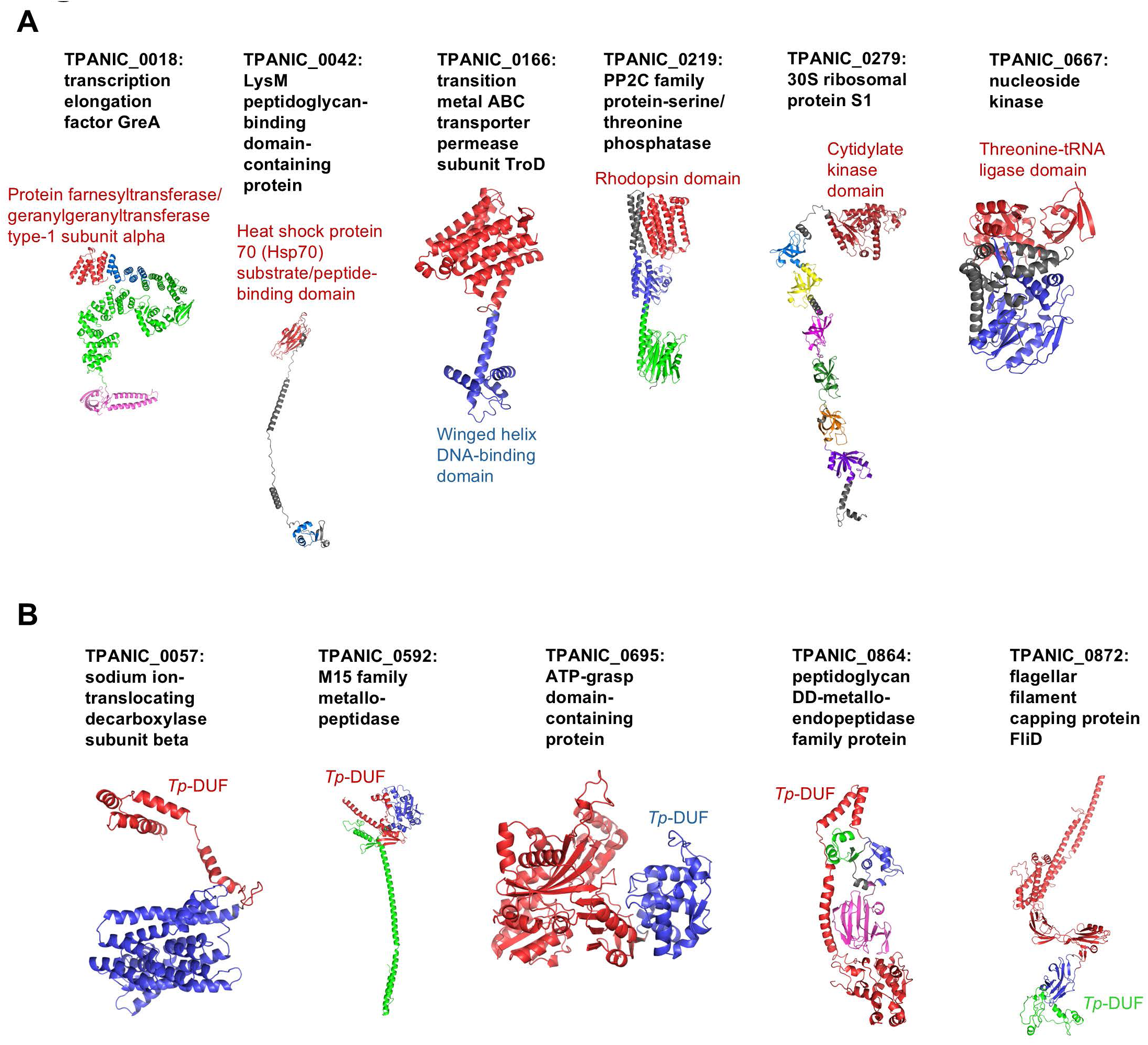
Structure models of *T. pallidum* proteins with predicted novel domain combinations and domains of unknown function. RoseTTAFold structure models of (A) six *T. pallidum* multi-domain proteins identified with potential novel domain combinations and (B) five *T. pallidum* multi-domain proteins with novel domains of unknown function (*Tp-*DUF). Individual domains are shown in different colours. Grey regions indicate predicted linker or unstructured regions between domains, or in the case of TPANIC_0667, regions that lacked confident DALI structural homology. Red regions in each ribbon diagram indicate the N-terminal domain. The protein locus tag and NCBI annotated protein functions are indicated above each protein model. The DALI-derived functional annotations of domains not observed in orthologous proteins are included in panel A, with colour-coding to match the corresponding ribbon diagram. Domains of unknown function are indicated by “*Tp*-DUF”.

### Structure-based functional annotation of *T. pallidum* proteins of unknown function

Among the 323 *T. pallidum* proteins of unknown functions, 240 were confidently modeled and assigned DALI-based functions via the identification of high-confidence structural ortholog matches (**Figure 1C and Tables S2 and S9_parts 1 and 2**), including most of the treponemal known and putative OMPs and several proteins identified as PPRPs, as described below. The 83 *T. pallidum* proteins of unknown function that could not be assigned DALI-based functional annotations failed to meet the minimum confidence cutoff scores for structure modeling and/or DALI comparisons, while a few failed due to protein size constraints or data processing errors (**Table S9_part 3**). These findings have allowed for the inference of function for the majority of proteins within a group of *T. pallidum* proteins that remain relatively uncharacterized despite comprising approximately one-third of the treponemal proteome. Further, these proteins are likely to be critical for the organism’s unique and successful mechanisms of pathogenesis.

### Identification of *T. pallidum* putative pathogenesis related proteins

Our AI-based structure-to-function workflow identified 26 high-confidence and high-ranking PPRPs, excluding known and putative OMPs (discussed separately below) (**Figure 1C and Tables 1 and S10**), seven of which were predicted in previous genomics and TBM studies (Cejkova et al. 2012; Houston et al. 2018; Petrosova et al. 2012) (**Table 1**). The PPRPs included eight functionally annotated proteins. These were predicted through AI-based modeling to be structural orthologs of bacterial proteins involved in infectivity, persistence, virulence regulation, OMP biogenesis, and macrophage invasion, survival, and replication (**Tables 1 and S10**). In these analyses, structural orthologs with a Z-score of 8.0 or higher were deemed to have sufficient structural homology to be considered possible orthologs of well-characterized bacterial proteins (as explained in the Methods section “Comparative structure analyses: DALI analyses and identification of high-confidence *T. pallidum* protein structure model orthologs”). These PPRPs included structural orthologs of the localization of lipoproteins (Lol) pathway, including LolB (TPANIC_0789; confidence score of Z=11.9), LolC (TPANIC_0580 or TPANIC_0582; confidence scores of Z=28.0 and Z=24.5), LolD (TPANIC_0581 or TPANIC_0964; confidence scores of Z=33.3 and Z=34.1), and LolE (TPANIC_0580 or TPANIC_0582; confidence scores of Z=27.9 and Z=23.6) (**Tables 1, S1, S2, and S10**). These proteins are involved in the transport of lipoproteins from the inner membrane to the outer membrane (Wilson and Bernstein 2016). The remaining 18 *T. pallidum* PPRPs were classified as proteins of unknown function by genome annotation. Almost half of these were identified by AI-based modeling as potential structural orthologs of proteins involved in host adhesion, protein/tissue destruction, and invasion. These included TprG (TPANIC_0317) and TprH (TPANIC_0610) from the 12-membered *T. pallidum* repeat (Tpr) protein family (LaFond and Lukehart 2006), which were predicted to be structural orthologs of known Gram-negative pathogenesis-related OMPs (**Tables 1 and S10**).

**Table 1.** *Treponema pallidum* PPRPs identified via AI-based structure modeling-to-function analyses.

| Locus Tag | DALI Structural Ortholog | Potential Roles in Virulence |
| --- | --- | --- |
| TPANIC_0086 | c-di-GMP receptor, PlzA (*Borrelia burgdorferi*) | Infectivity and persistence |
| TPANIC_0110 | Encapsulin cargo/iron-binding protein (*Bacillus thermotolerans*) | Iron sequestration, storage, and homeostasis |
| TPANIC_0183 | TolB (*Escherichia coli*) | Maintenance of outer membrane integrity |
| TPANIC_0225<sup>†</sup> | Choline-binding protein, PcpA (*Streptococcus pneumoniae*) | Host adhesion |
| TPANIC_0262<sup>†</sup> | cAMP-responsive global transcriptional regulator, MtbCRP (*Mycobacterium tuberculosis*) | Global transcriptional regulator of virulence |
| TPANIC_RS01325 | Tetanus and botulinum neurotoxins (C-terminal region of heavy chain), TeNT and BoNT (*Clostridium botulinum*, *Clostridium tetani*) | Toxin binding/translocation into neuronal cells |
| TPANIC_0317<sup>†</sup> | Outer membrane protein, PorB (*Neisseria meningitidis*) | Immune evasion and adhesion/invasion of host epithelial cells |
| TPANIC_0369 | Beta-barrel assembly machinery protein, BamD (*Rhodothermus marinus*) | Essential protein for OMP biogenesis |
| TPANIC_0455 | Extracellular toxic zinc metalloendopeptidase, AsaP1 (*Aeromonas salmonicida*) | Host protein degradation/tissue damage (necrosis, ulceration, tissue lysis) |
| TPANIC_0473 | Mycobacterial ESX-3 translocon protein/Type VII secretion system protein, EccD3 (*Mycobacterium smegmatis*) | Virulence-related protein export |
| TPANIC_0580 | Lipoprotein outer membrane localization protein C/E, LolC/LolE (*Escherichia coli*) | Lipoprotein transport from inner to outer membrane |
| TPANIC_0581 | Lipoprotein outer membrane localization protein D, LolD (*Aquifex aeolicus*) | Lipoprotein transport from inner to outer membrane |
| TPANIC_0582 | Lipoprotein outer membrane localization protein C/E, LolC,/LolE (*Escherichia coli*) | Lipoprotein transport from inner to outer membrane |
| TPANIC_0610<sup>†</sup> | Outer membrane protein/protease, OmpT (*Vibrio cholerae*) | Host protein degradation and outer membrane vesicle-mediated toxin delivery |
| TPANIC_0693 | Outer membrane inner leaflet localized lipoprotein, PelC (*Geobacter metallireducens*) | Exopolysaccharide secretion/export |
| TPANIC_0768 | Lyme disease major virulence factor, BB0323 (*Borrelia burgdorferi*) | Infectivity and outer membrane integrity |
| TPANIC_0789<sup>†</sup> | Outer membrane lipoprotein, LolB | Lipoprotein transport from inner to outer membrane |
| TPANIC_0833 | Putative adhesin (*Bacteroides* and *Clostridium* sp.) | Host adhesion |
| TPANIC_0839 | Putative adhesin (*Bacteroides* and *Parabacteroides* sp.) | Host adhesion |
| TPANIC_0862<sup>†</sup> | Macrophage infectivity potentiator protein, Mip (*Legionella pneumophila*) | Macrophage invasion/survival/replication and host tissue/extracellular matrix adhesion |
| TPANIC_0930 | Glycosyltransferase/Motility associated factor, Maf (*Magnetospirillum magneticum*) | Flagella O-glycosylation, function, and motility |
| TPANIC_0962 | ABC transporter/macrolide export ATP-binding/permease protein, MacB (*Acinetobacter baumannii*) | Antibiotic resistance, and virulence factor export |
| TPANIC_0963 | ABC transporter/macrolide export ATP-binding/permease protein, MacB (*Acinetobacter baumannii*) | Antibiotic resistance, and virulence factor export |
| TPANIC_0964 | Lipoprotein outer membrane localization protein D, LolD (*Aquifex aeolicus*)<br>ABC domain, ABC transporter/macrolide export ATP- | Lipoprotein transport from inner to outer membrane<br>Antibiotic resistance, and virulence factor export |
|  | binding/permease protein, MacB<br>(*Escherichia coli*) |  |
| TPANIC_0965 | Membrane fusion protein/macrolide-specific efflux protein, MacA<br>(*Escherichia coli*) | Antibiotic resistance, and virulence factor export |
| TPANIC_1033<sup>†</sup> | Type V secreted phospholipase, PlpD<br>(*Pseudomonas aeruginosa*) | Tissue invasion and immune evasion. |

<sup>†</sup>Seven proteins identified as PPRPs in previous genomics and template-based modeling studies (Cejkova et al. 2012; Houston et al. 2018; Petrosova et al. 2012).

The fibronectin type III (Fn III) domain-containing protein TPANIC_RS01325 was identified as a high-confidence structural ortholog of the C-terminal region of the heavy chain (Hc) of clostridial neurotoxins, TeNT (Fotinou et al. 2001) and BoNT (Karalewitz et al. 2012), with most aligned residues located outside the Fn III domain (**Tables 1 and S10 and Figure S1**).

TPANIC_RS01325 also showed structural similarity to nontoxic nonhemagglutinin (NTNH) (**Tables S2 and S10**), a structural ortholog of clostridial neurotoxins that stabilizes and protects them from degradation (Gu et al. 2012). Also of interest, TPANIC_0262, a potential structural ortholog of the global virulence regulator MtbCRP in *Mycobacterium tuberculosis*, is located immediately upstream of TPANIC_RS01325 in the proteome (**Tables 1 and S10**). Most of the remaining PPRPs were identified as potential structural orthologs of proteins involved in the secretion and export of molecules across the cell envelope and/or into the external host environment (**Tables 1 and S10**). DeepLocPro (Moreno et al. 2024) predictions suggest that at least nine PPRPs may be secreted into the extracellular environment or localized to the outer membrane of *T. pallidum* (**Table S10_part 2**). Together, these results offer AI-based insights into *T. pallidum* pathogenesis via the identification of treponemal PPRPs.

### Bioinformatic and structural analysis of known and previously predicted *T. pallidum* OMPs

Using AlphaFold 3 modeling, we previously showed that 19 known and potential *T. pallidum* OMPs exhibit Nichols-SS14 inter-strain amino acid sequence variability, with most variant residues primarily located in surface-exposed, host-facing regions of the proteins (Houston et al. 2024b). More recently, two studies using the same AlphaFold 3 structural mapping approach supported our earlier findings by demonstrating that inter-strain variable amino acids in seven OMPs from *T. pallidum* clinical strains are also primarily located in surface-exposed regions (Bettin et al. 2025; Pospisilova et al. 2025). In the present study, we used multi-method AI-based structure modeling with RoseTTAFold and AlphaFold 3, in combination with DALI structure comparison analyses and sequence-based computational tools, to analyze the predicted structure, function, and subcellular localization of 36 known and predicted *T. pallidum* OMPs (Anand et al. 2015; Anand et al. 2012; Brinkman et al. 2008; Cameron 2003; Cameron et al. 2004; Cameron et al. 2000; Centurion-Lara et al. 1999; Centurion-Lara et al. 2013; Cox et al. 2010; Desrosiers et al. 2011; Djokic et al. 2019; Giacani et al. 2015; Giacani et al. 2005; Hardham and Stamm 1994; Hawley et al. 2021; Haynes et al. 2021; Houston et al. 2012; Houston et al. 2018; McGill et al. 2010; Osbak et al. 2016; Parker et al. 2016; Radolf and Kumar 2017) (**Figure 1C**). As shown in **Figures S2-S6 and Table S11_part 1**, 35 OMPs were confidently modeled by at least one AI-based program, and 31 were modeled with high-confidence by both programs. OMP surface mapping revealed minor differences between the two modeling programs, notably in the beta-barrel/extracellular loop (ECL) interfaces (**Figures S2-S5**). Therefore, we used the AI-based structure modeling predictions to generate OMP ECL consensus sequences (**Table S11_part 1**) to aid in the identification of surface-exposed BCEs, as discussed below.

Structural differences of four TolC-like OMPs from *T. pallidum* (TPANIC_0966-TPANIC_0969) were observed between the AI-based models in the present study, previous TBM predictions (Hawley et al. 2021; Houston et al. 2018; Radolf and Kumar 2017), and the crystal structure of *Escherichia coli* TolC (Higgins et al. 2004). In the TBM studies, TolC models predicted four beta-strands per monomer that trimerize to form a 12-stranded beta-barrel pore with six ECLs, a structure similar to the crystal structure solved for *E. coli* TolC (Hawley et al. 2021; Higgins et al. 2004; Houston et al. 2018; Radolf and Kumar 2017) (**Figure S7A and S7B**). Both AI-based modeling programs in the present study predicted eight-strands per monomer that form 24-stranded beta-barrel pores with 12 ECLs upon trimerization (**Figure S7A and S7B**). The AlphaFold 3 multimer also predicted larger beta-barrel pore channels in each of the four treponemal proteins (∼45A diameter) compared to *E.coli* TolC (∼30A diameter) (Higgins et al. 2004) (**Figure S7C**).

Although 31 of the 36 known and potential OMPs lack clear functional annotations in the NCBI proteome, 35 were assigned confident predicted functions based on DALI structural comparisons in the present study (**Tables 2 and S11_part 2**). These included OMPs/porins involved in the transport and uptake of nutrients, in particular iron, vitamins, antibiotics, and small molecules e.g. organic acids and oligosaccharides (**Tables 2 and S11_part 2**). Thirty of the known or potential *T. pallidum* OMPs were identified as PPRPs, including nine of the 12-membered *T. pallidum* repeat (Tpr) protein family (LaFond and Lukehart 2006) and 13 OMPs not previously listed as PPRPs in prior genomics and TBM studies (Cejkova et al. 2012; Houston et al. 2018; Petrosova et al. 2012) (**Tables 2 and S11_part 2**). Several of the newly-identified PPRPs were predicted to have functions related to immune evasion, host adhesion and invasion, and protein export/secretion, including TPANIC_0483, which had previously been experimentally validated as a treponemal adhesin (Cameron et al. 2004) (**Tables 2 and S11_part 2**). Of the five OMPs with assigned functions in the NCBI annotated proteome, four matched our structure-to-function annotation (**Tables 2 and S11_part 2**). Among the 36 known or potential OMPs, eight were identified as having structural folds lacking typical Gram-negative-like integral OMP beta-barrel domains and high-confidence structural ortholog matches not typically associated with outer membrane localization (**Tables 2 and S11_part 2**). Of note, TPANIC_0136 and TPANIC_0751 have been identified as lipoproteins, with experimental evidence supporting TPANIC_0136 binding extracellular matrix (ECM) components and epithelial and glioma cells (Brinkman et al. 2008; Djokic et al. 2019; Ke et al. 2015). Similarly, TPANIC_0751 is a well-characterized vascular adhesin that has been shown to bind ECM components and the endothelial receptor LamR that is bound by other neuroinvasive pathogens (Cameron 2003; Houston et al. 2011; Houston et al. 2012; Kao et al. 2017; Lithgow et al. 2020; Parker et al. 2016).

Previous experimental findings, together with structure modeling, DALI analyses, and bioinformatics data from the present study, were used to classify the 36 proteins into OMP confidence categories. Proteins were classified as high-confidence OMPs if previous experimental data demonstrated outer membrane localization (Anand et al. 2012; Brinkman et al. 2008; Cameron et al. 2000; Centurion-Lara et al. 1999; Desrosiers et al. 2011; Houston et al. 2012; Parker et al. 2016). In the absence of prior experimental data, proteins were assigned to the high-confidence category based on the following criteria: (1) at least half of the algorithms predicted the presence of a cleavable signal peptide; (2) at least one of the algorithms predicted outer membrane localization; (3) at least one of the sequence-based beta-barrel prediction algorithms predicted the presence of a beta-barrel structure comprised of at least eight beta-strands; (4) AI-based structure modeling predicted the presence of a beta-barrel (unless prior experimental evidence already confirmed outer membrane localization); and (5) DALI analyses identified top-ranking, high-confidence structural orthologs that are known outer membrane-localized proteins from Gram-negative bacteria. Proteins that did not meet all the high-confidence criteria were classified as either medium- or low-confidence OMPs, depending on the number and nature of the unmet criteria. As shown in **Tables S11_part 3 and S12**, 25 proteins were categorized in the higher OMP confidence category, four proteins were categorized as medium confidence, and seven proteins were classified as low-confidence OMPs. Together, these findings update the bioinformatics-based structural and functional characterizations of previously identified and predicted *T. pallidum* OMPs.

### Structure modeling-based identification and bioinformatic characterization of novel *T. pallidum* OMP candidates

AI-based modeling and DALI analyses identified seven *T. pallidum* proteins of unknown function with previously unreported beta-barrel domains that are structurally similar to known OMPs from other bacterial species (**Figures 1C and 6 and Table S13**). Previous MS-based proteomics studies have shown that at least five of these proteins are expressed in *T. pallidum* (Houston et al. 2023; Houston et al. 2024a; Osbak et al. 2016) (**Table S13_part 1**). With beta-barrel domain folds common in surface-exposed, host-facing OMPs in Gram-negative bacteria (Doyle and Bernstein 2024), we evaluated these seven treponemal proteins as potential novel OMP candidates. Highly similar tertiary folds and beta-barrel domains containing at least eight beta-strands were predicted by both AI-based modeling programs in six of the seven proteins; the only exception was TPANIC_RS05525, in which the AlphaFold monomer was modeled as a 10-stranded open beta-sheet-containing protein (**Figure 6**). Notably, AlphaFold 3 multimerization analyses generated a confident model for the dimer of TPANIC_RS05525 (**Figure 6**).

**Figure 6.**
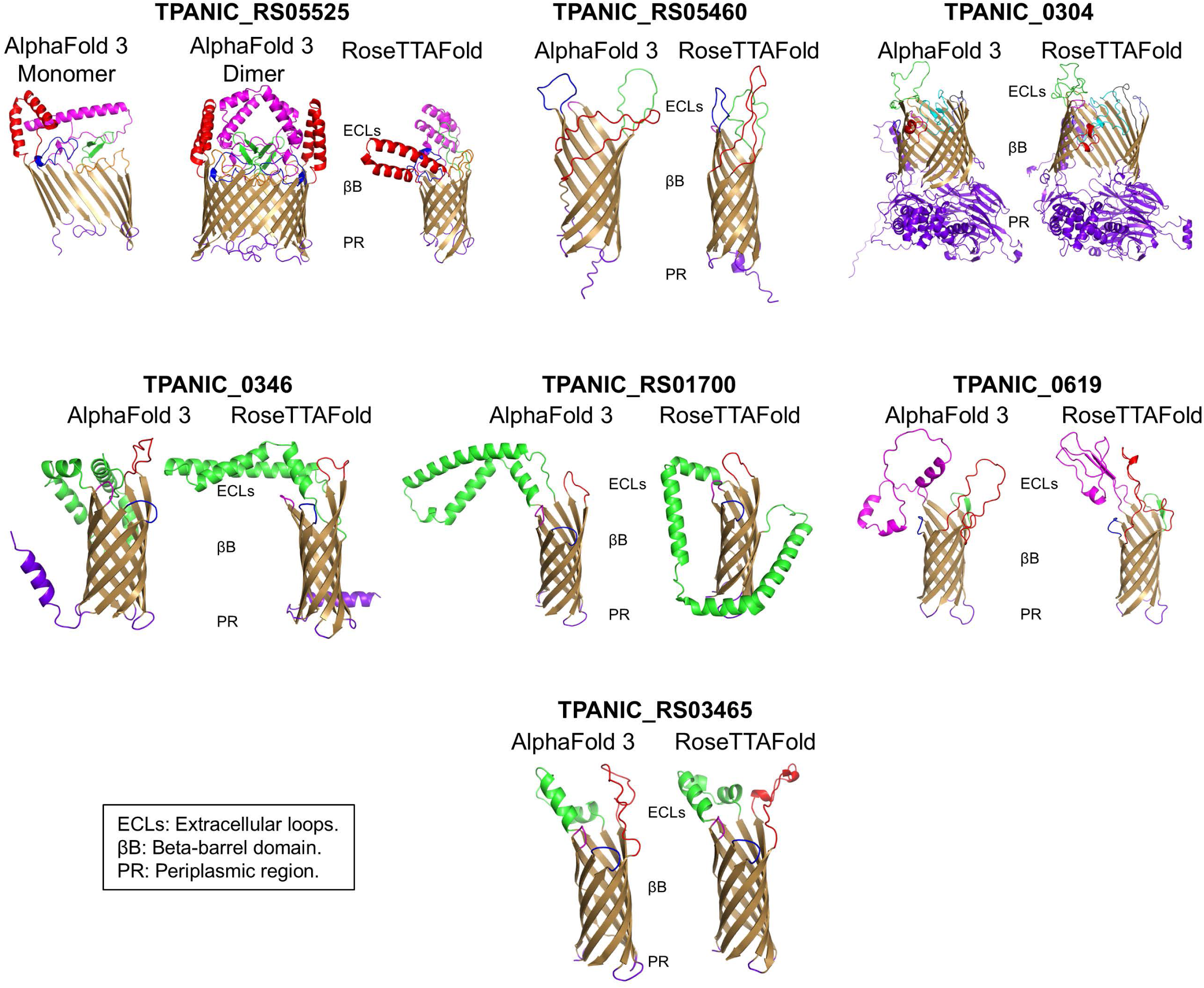
AI-based structure modeling of seven potential novel *T. pallidum* OMPs. AlphaFold 3 and RoseTTAFold models showing lateral views of seven *T. pallidum* proteins predicted to be OMPs. These proteins have not been previously identified as potential OMPs. The monomeric and dimeric models of TPANIC_RS05525 are shown; dimer models were generated using AlphaFold 3-Multimer. Extracellular loops (ECLs) are highlighted in different colours and are orientated at the top of each beta barrel (surface-exposed/host-facing region). Predicted integral outer membrane beta-barrel (βB) domains are shown in bronze. Predicted periplasmic-localized protein regions (PR) are shown in purple (base of the beta-barrel domains).

Additionally, DALI analyses of the full-length TPANIC_0304 protein model focused only on the N-terminal region of TPANIC_0304, which was not predicted to contain a beta-barrel domain in the present study (**Table S2 and Figure 6**). However, additional DALI analyses focused on the TPANIC_0304 C-terminal region identified high-confidence structural orthologs corresponding to beta-barrel domains of established OMPs from Gram-negative bacteria (**Table S13_parts 2 and 3**).

Using our previously described inter-strain variable amino acid comparative analyses and AI-based structural mapping approach (Houston et al. 2024b), we found that four of the seven potential novel OMPs are annotated with Nichols-SS14 inter-strain variable residues. Of these four proteins, three were found to exhibit sequence variations in predicted surface-exposed, host facing regions and ECLs: TPANIC_RS05525, TPANIC_0304, and TPANIC_0346 (**Figure S8 and Table S13_part 4**). Data integration analyses, which incorporated MS experimental data from previous studies (Houston et al. 2023; Houston et al. 2024a; Houston et al. 2024b; Osbak et al. 2016; Romeis et al. 2021), confirmed the presence of annotated inter-strain protein sequence differences in two of the potential novel OMPs: TPANIC_0304, and TPANIC_0346 (**Figure S8 and Tables S13_part 4**). ECL consensus sequences were generated for the seven proteins, as described previously (**Table S13_part 1**).

DALI-based analyses assigned putative functions to all seven potential novel OMPs (**Tables 3 and S13_parts 2 and 3**). Furthermore, all seven potential novel OMPs were identified as PPRPs (**Tables 3 and S13_parts 2 and 3**), bringing the total number of *T. pallidum* PPRPs to 63, including both known/potential OMPs (**Tables 2, 3, S11_part 2 and S13_parts 2 and 3**) and non-OMPs (**Tables 1 and S10**). Predicted functions included nutrient uptake/transport, outer membrane structural integrity/biogenesis, host adhesion/invasion, immune modulation/evasion, autotransporter/toxin secretion, and antibiotic/antimicrobial peptide (AMP) resistance (**Tables 3 and S13_parts 2 and 3**).

**Table 2.**
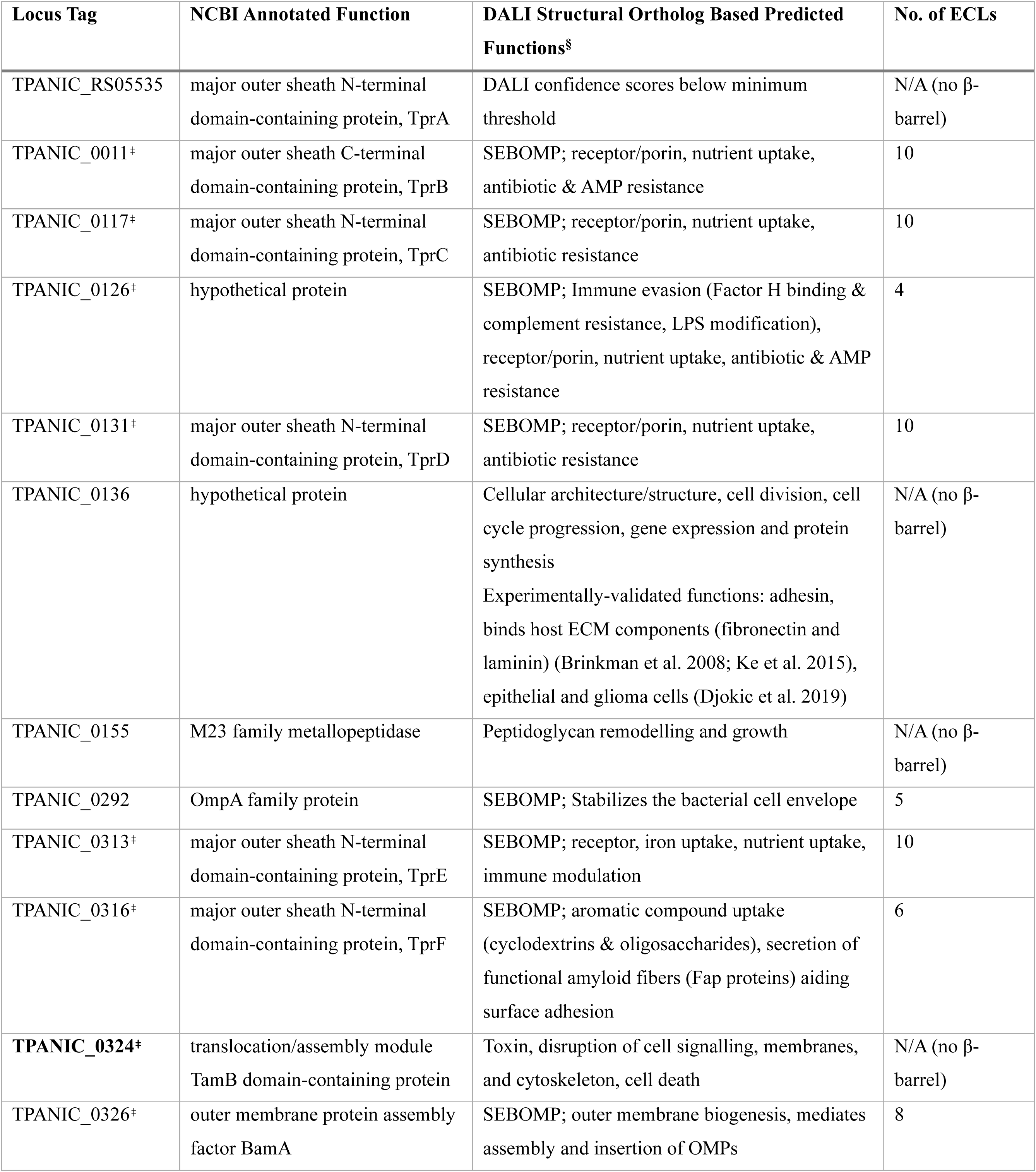
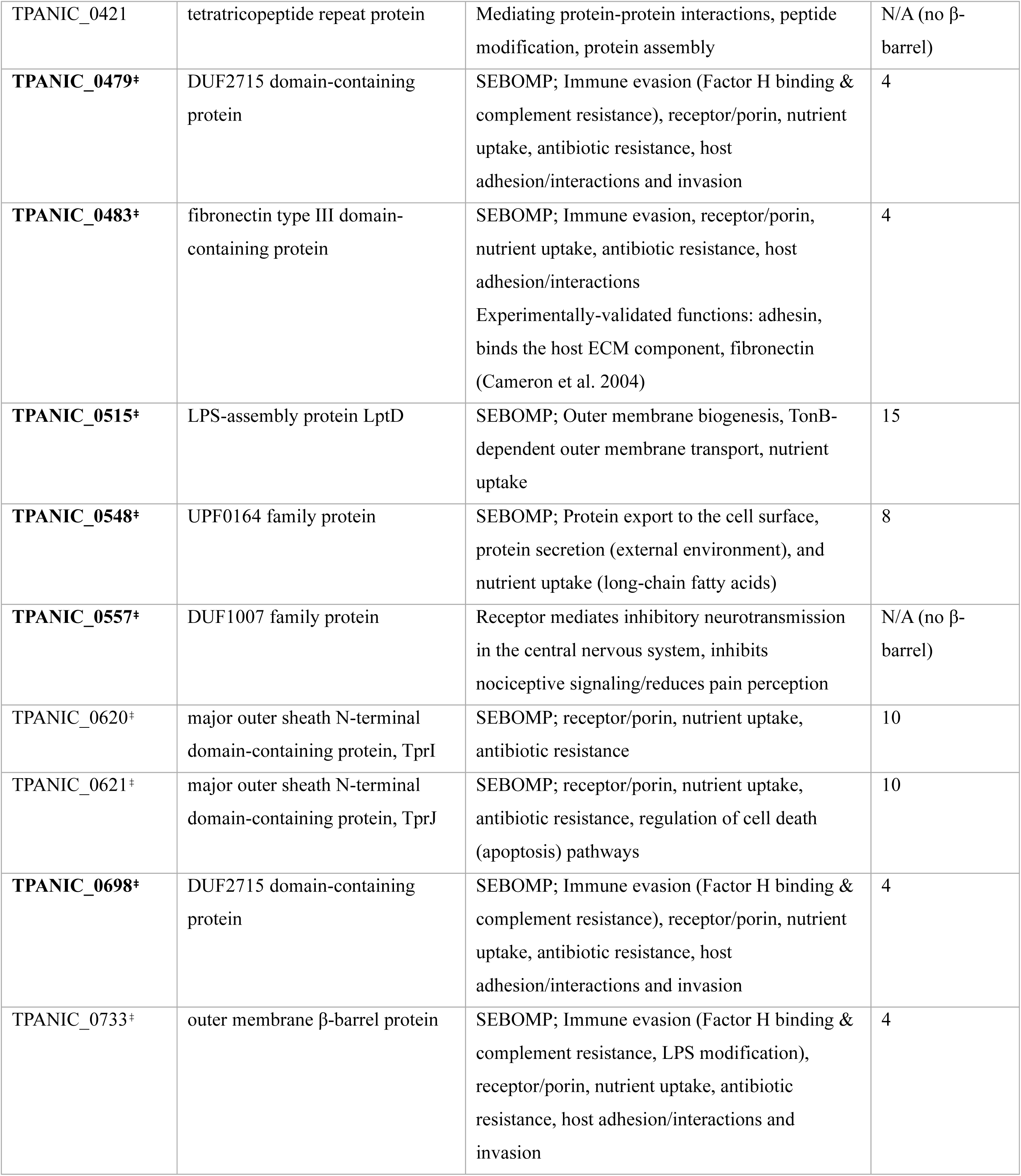
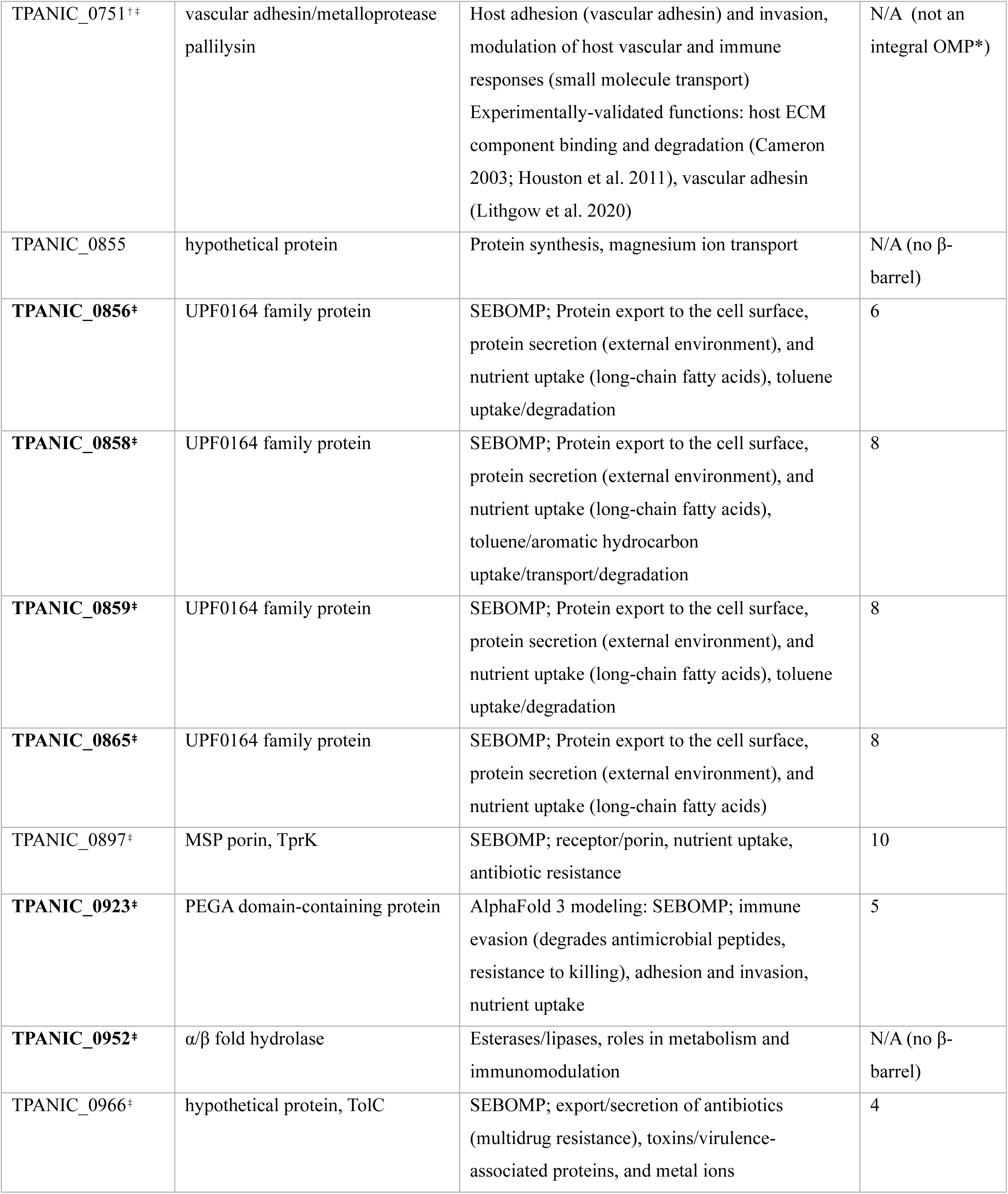
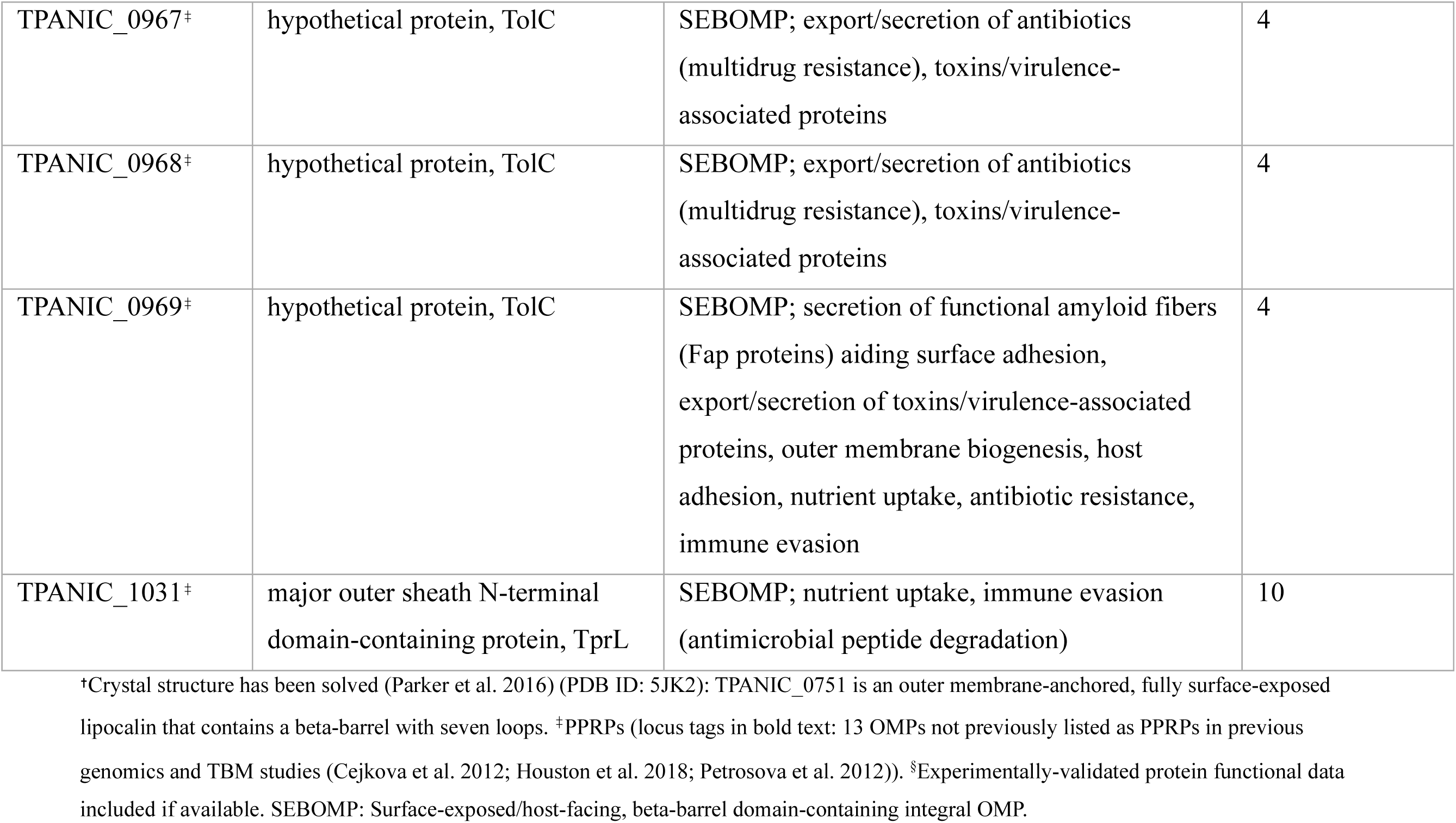
AI-based structural and functional characterization of 36 known and predicted *T. pallidum* OMPs.

**Table 3.** AI-based structural and functional characterization of seven potential novel *T. pallidum* OMPs.

| Locus Tag | NCBI Annotated Function | DALI Structural Ortholog Based Predicted Functions | No. of ECLs |
| --- | --- | --- | --- |
| TPANIC_RS05525<sup>†</sup> | major outer sheath C-terminal domain-containing protein | SEBOMP; host adhesion/interactions, host tissue degradation and invasion, immune evasion, AMP degradation/resistance, receptor/porin, nutrient uptake | 5 |
| TPANIC_RS05460<sup>†</sup> | hypothetical protein | SEBOMP; outer membrane structural integrity, host adhesion, immune modulation and evasion, receptor/porin, nutrient uptake | 4 |
| TPANIC_0304<sup>†</sup> | hypothetical protein | SEBOMP; outer membrane integrity and biogenesis, mediates assembly and insertion of OMPs, receptor/porin, nutrient uptake, autotransporter and toxin secretion, inhibition & killing of competing microbes | 8 |
| TPANIC_0346<sup>†</sup> | DUF2715 domain-containing protein | SEBOMP; Immune modulation and evasion (including complement and serum resistance), host adhesion and invasion, receptor/porin, nutrient uptake, antimicrobial resistance, antibiotic resistance | 4 |
| TPANIC_RS01700<sup>†</sup> | DUF2715 domain-containing protein | SEBOMP; outer membrane structural integrity and stability, receptor/porin, nutrient uptake, AMP resistance, host adhesion and invasion, antibiotic resistance, immune evasion (resistance to complement-mediated killing, serum resistance) | 4 |
| TPANIC_0619<sup>†</sup> | DUF2715 domain-containing protein | SEBOMP; receptor/porin, nutrient uptake, AMP resistance, immune evasion (complement activation inhibition, immune modulation), outer membrane structural integrity and stability, host adhesion and invasion | 4 |
| TPANIC_RS03465<sup>†</sup> | DUF2715 domain-containing protein | SEBOMP; outer membrane structural integrity and stability, receptor/porin, nutrient uptake, host adhesion and invasion, immune evasion (resistance to complement-mediated killing, surface molecule modification), antibiotic resistance, AMP resistance | 4 |

<sup>†</sup>PPRPs (all seven predicted OMPs not previously listed as PPRPs in previous genomics and TBM studies (Cejkova et al. 2012; Houston et al. 2018; Petrosova et al. 2012)). SEBOMP: Surface-exposed/host-facing, beta-barrel domain-containing integral OMP.

Using our OMP confidence classification criteria, two of the potential novel OMPs were ranked as high-confidence and five as medium-confidence (**Tables S13_part 5 and S14**). Notably, the two high-confidence novel OMP candidates, TPANIC_RS03465 and TPANIC_0346, are located directly adjacent to other OMP candidates in the *T. pallidum* proteome. Several of the other novel OMP candidates also cluster near Tpr family OMPs or within OMP-rich regions of the proteome (data not shown). When combined, the findings from the present study enabled the compilation of a refined list of 36 *T. pallidum* proteins, comprising six known OMPs (Anand et al. 2012; Brinkman et al. 2008; Cameron et al. 2000; Centurion-Lara et al. 1999; Desrosiers et al. 2011; Houston et al. 2012; Parker et al. 2016) and 30 high- to medium-confidence candidate OMPs. Of these 30 candidates, 21 were classified as high-confidence and nine as medium-confidence OMPs (**Table S15**). This updated list of previously identified and newly predicted *T. pallidum* OMPs provides a resource for prioritizing treponemal protein targets in syphilis vaccine design.

### Identification of potential *T. pallidum* linear B cell epitopes

Multi-server prediction analyses identified 2680 putative linear BCEs across the 36 proteins from the refined list of known and potential OMPs, including overlapping epitopes (**Figure 1C and Table S16_part 1**). The 2680 predicted BCEs were mapped onto the structure models of the 36 known and potential OMPs, which identified 1133 potential BCEs that contained at least five surface-exposed, host-facing amino acids (**Table S16_part 2**). These BCEs were further prioritized using eight additional criteria based on bioinformatic analyses, biophysical properties, amino acid sequence conservation, and previously reported protein expression data. Among the surface-exposed *T. pallidum* BCE candidates, 103 fulfilled all nine criteria and corresponded to 22 known and potential OMPs (**Table S16_parts 3 and 4**). Of these, 92 putative BCEs were conserved across all *T. pallidum* strains but differed from one another by at least one amino acid residue (**Table S17 and Figure S9**); eighteen of these BCEs were identified in a previous TBM study (Hawley et al. 2021), while 74 were predicted for the first time in the present analyses (**Table S17**). Each of these 92 BCEs mapped to either two proteins with experimental evidence supporting their OMP annotations (TPANIC_0751 and TPANIC_0897) (Centurion-Lara et al. 1999; Houston et al. 2012; Parker et al. 2016), or to 21 candidate OMPs including five of the seven potential novel OMPs identified in the present study (**Figure S9 and Table S17**).

## Discussion

Although TBM programs have been used to model *T. pallidum* proteins (Hawley et al. 2021; Radolf and Kumar 2017), including the whole proteome (Houston et al. 2018), and AlphaFold 3 has been applied to a subset of *T. pallidum* proteins (Bettin et al. 2025; Delgado et al. 2024; Houston et al. 2024b; Pospisilova et al. 2025), comprehensive AI-based structure modeling analyses of the *T. pallidum* proteome have not been previously performed. In this work, we applied the deep learning modeling programs, RoseTTAFold (Baek et al. 2021) and AlphaFold (Abramson et al. 2024), within a global structure modeling-to-function pipeline. Method validation analyses revealed a strong correlation between protein functional annotations generated from the NCBI Prokaryotic Genome Annotation Pipeline (PGAP) (Li et al. 2021; Tatusova et al. 2016) and our AI-based modeling pipeline, supporting the validity of our workflow. This validated structure modeling-to-function pipeline enabled the generation of high-confidence three-dimensional models and structure-based functional annotations for almost 90% of the proteins in the *T. pallidum* proteome. It also provides a broadly applicable approach for the computational, system-level characterization of proteins and their domains from genetically intractable and uncultivable pathogens, including other spirochetes and fastidious bacteria of global health importance.

The concept of domain consolidation, where domain architectures that are usually found across multiple separate proteins in other species are merged into single proteins, is a hallmark of genome reduction that often results in increased protein functional complexity (Kelkar and Ochman 2013). Here, we identified six treponemal proteins with domain combinations not previously reported in orthologous proteins from other species, based upon literature searches. These included TPANIC_0279, a 30S ribosomal S1 protein that also contains a cytidylate kinase domain. Another unique domain architecture was observed for the LysM peptidoglycan-binding protein, TPANIC_0042, which was predicted to contain an N-terminal heat shock protein 70 (Hsp70) substrate/peptide-binding domain. Finally, the transition metal ABC transporter permease subunit TroD protein, TPANIC_0166, was predicted to contain a C-terminal winged helix DNA-binding domain that has been shown to play a role in transcriptional regulation (Desrosiers et al. 2010; Pohl et al. 1999). These findings contribute to the functional annotation of *T. pallidum* by the identification of potential multi-functional proteins that are not fully annotated in proteome databases. They also support the concept that domain consolidation is an evolutionary strategy that maximizes functional efficiency and versatility in the highly reduced genome of this obligate human pathogen. We also identified five *T. pallidum* proteins with at least one modeled domain that lacked DALI structural orthologs. Since protein tertiary structure is a key determinant of protein function, these domains of unknown function may mediate novel or previously unidentified functions in these proteins.

Our previous TBM study demonstrated the potential effectiveness of protein structure modeling for the prediction of protein structures and functions on a global scale (Houston et al. 2018). However, since TBM programs rely on the availability of protein structures with high levels of sequence homology (Kelley et al. 2015), their effectiveness in generating accurate models decreases in the absence of such structures (Baker and Sali 2001). As a result, in the prior TBM-based studies only partial models were generated for the majority of the *T. pallidum* proteins, and 20% of the *T. pallidum* proteome could not be modeled with high-confidence or assigned predicted functions (Houston et al. 2018). Notably, the non-modeled proteins included half of the proteins of unknown function, due to the lack of relevant protein structures (Houston et al. 2018). To overcome these limitations and to improve protein modeling accuracy, we used a combination of AI-based structure modeling and comparative structure analyses in the present study. This approach provided insight into mechanisms of *T. pallidum* pathogenesis by the generation of high-confidence models with 100% sequence coverage for nearly all treponemal proteins. This structure-to-function workflow allowed for the identification of *T. pallidum* proteins that are high-confidence structural orthologs of LolC, LolD, and LolE. TPANIC_0581 was identified as a high-confidence structural ortholog of LolD; additionally, TPANIC_0964 was identified as another potential structural ortholog of LolD, and also had structural similarity to an ABC transporter/macrolide export ATP-binding/permease protein. TPANIC_0580 and TPANIC_0582 were both structural orthologs of both LolC and LolE. These proteins comprise an inner membrane ATP-binding cassette (ABC) transporter complex that extracts lipoproteins from the inner membrane and transfers them to the periplasmic carrier protein, LolA (annotated in the proteome as TPANIC_0333, consistent with the structure modeling in the present study). In Gram-negative bacteria, LolA transports the lipoprotein to LolB for insertion into the outer membrane (Wilson and Bernstein 2016). In *T. pallidum*, structural modeling of TPANIC_0789 showed a structural ortholog match to LolB with a Z-score in the significant range. Together, these findings suggest that *T. pallidum* may possess a conserved Lol pathway, a bioinformatic prediction that should be followed up with wet lab-based experimental validation.

One of the main aims of the present study was to identify putative functions for the large subset of *T. pallidum* proteins of unknown function, many of which may contribute to treponemal survival and pathogenesis. In the absence of significant amino acid sequence homology, our structure-based modeling approach enabled high-confidence functional predictions for three-quarters of the *T. pallidum* proteins of unknown function. In addition to proteins involved in essential “housekeeping” processes, we identified putative functions for proteins with potential roles in *T. pallidum* pathogenesis. These included most of the known and potential integral OMPs.

Surface-exposed *T. pallidum* OMPs have been identified as leading syphilis vaccine candidates due to their location at the bacterial-host interface and their recognition by protective antibodies during infection (Delgado et al. 2022; Giacani et al. 2010; Haynes et al. 2021; LaFond and Lukehart 2006; Lithgow et al. 2017; Waugh and Cameron 2024). In the current study, we compiled an up-to-date, refined list of 36 known and potential *T. pallidum* OMPs. Of these, six were experimentally confirmed OMPs (Anand et al. 2012; Brinkman et al. 2008; Cameron et al. 2000; Centurion-Lara et al. 1999; Desrosiers et al. 2011; Houston et al. 2012; Parker et al. 2016) and 30 were high- to medium-confidence candidates, including seven newly identified potential OMPs. In a previous study by Cox et al. that used amino acid sequence-based bioinformatics tools to rank 19 OMP candidates into four confidence categories (Cox et al. 2010), the six highest-ranking proteins were also identified as high-confidence OMPs in the present study.

Additionally, half of the lowest-ranking candidates from the previous study (Cox et al. 2010) were not predicted to be OMPs in the present study. More recently, a TBM study defined the *T. pallidum* outer membrane protein repertoire (the “OMPeome”) as comprising 26 OMPs (Hawley et al. 2021). In the present study, 25 of these 26 proteins were also identified as known and potential OMPs, and the OMP repertoire was expanded by 11 additional candidates; seven novel OMPs identified in the current study, as well as TPANIC_0292, TPANIC_0483, and two experimentally validated OMPs, TPANIC_0136 (Brinkman et al. 2008) and TPANIC_0751 (Houston et al. 2012; Parker et al. 2016). The identification of novel *T. pallidum* OMP candidates in the present study provides new targets for syphilis protein subunit vaccine development.

Identifying treponemal peptides that contain B cell epitopes is a key step in the design and development of syphilis subunit vaccines. Using BCE predictions and AlphaFold 3 modeling, a recent study identified several predicted epitopes in variable and conserved surface-exposed regions of six *T. pallidum* known and predicted OMPs (Bettin et al. 2025). In the present study, our AI-based modeling approach identified 1133 potential surface-exposed BCEs across all 36 known and predicted *T. pallidum* OMPs. Subsequent filtering based on established BCE criteria identified approximately 100 high-priority BCEs across 22 known and potential OMPs, including five of the seven potential novel OMPs identified in the present study. Most of the high-priority predicted BCEs had not been previously reported, reflecting the fact that our study analyzed the complete, expanded OMP repertoire, and included potential novel OMPs that have not been previously analyzed for BCEs.

The present structure-to-function modeling study has certain limitations. First, the DALI PDB searches depend on the availability of high-resolution protein structures in the PDB that are close structural orthologs of the *T. pallidum* query models. In cases where such homologous structures are absent, confident structure-based functional inference is not possible. However, only ∼10% of models could not be assigned functions due to the lack of high-confidence structural ortholog matches in the DALI searches. Second, structure-based functional inference was complicated when multiple structural orthologs with unrelated functions, but similar confidence scores, were generated in the DALI searches for a single *T. pallidum* protein model. However, this limitation affected less than 2% of all functionally annotated proteins in the present study. Third, proteins of unknown function were modeled and assigned putative functions with lower confidence compared to functionally annotated proteins, which may reduce the reliability of the assignment of predicted functions. Fourth, structure models were primarily generated for the Nichols strain of *T. pallidum*. Inter-strain amino acid sequence variations may alter the tertiary structures of orthologs in other *T. pallidum* strains, potentially affecting the composition of ECLs and the location of predicted BCEs. Finally, this structure-to-function approach is based entirely on modeling and bioinformatics-based predictions, and lacks experimental validation. For example, although approximately 100 high-priority BCE candidates were identified, an additional 1030 predicted epitopes did not meet all nine BCE prediction criteria. Some of these may correspond to more biologically relevant BCEs not captured by the strict computational thresholds. Ultimately, any epitope predictions will require experimental validation.

## Conclusion

The current study presents the first AI-based structure modeling analysis of the complete *T. pallidum* proteome. This computational approach has provided new insights into treponemal protein domain architectures, predicted functions, putative pathogenesis related proteins, known and potential OMPs, and candidate BCEs. Together, these findings offer a framework for advancing understanding of *T. pallidum* pathogenesis and informing syphilis vaccine development via the identification of novel target proteins and epitopes in globally circulating strains of *T. pallidum*.

## Materials and Methods

### Dataset: Treponema pallidum ssp. pallidum proteome

The *T. pallidum* ssp. *pallidum* Nichols strain proteome (NCBI Reference Sequence: NC_021490.2, annotated 12^th^ March 2023) was used for proteome-wide protein structure modeling, structure-to-function based protein annotations, and multi-domain analyses (**Figure 1A**). The proteome contains 980 proteins from predicted protein-coding genes, and seven proteins potentially encoded by “pseudo genes”. Additionally, three proteins that were previously shown to be expressed by *T. pallidum* using mass spectrometry, but were incorrectly deleted from the proteome, were included: TPANIC_0126a (Houston et al. 2024a; Osbak et al. 2016), TPANIC_0415 (Houston et al. 2024a), and TPANIC_RS05180 (Houston et al. 2024a). The amino acid sequence for TPANIC_RS04705 (NCBI proteome annotation, August 2019) was used for modeling instead of the TPANIC_RS05630 sequence (revised NCBI proteome annotation, March 2023), as previous mass spectrometry data indicated that TPANIC_RS05630 has an incorrectly truncated N-terminus, whereas TPANIC_RS04705 includes the correct extended N-terminal region (Houston et al. 2023). Of these 990 proteins, 667 were identified as proteins with assigned functional annotations or experimentally-determined functions, and 323 were identified as proteins of unknown function (“hypothetical proteins”, domain of unknown function [DUF] proteins, and proteins with limited or non-specific functional annotations, e.g. RNA-binding protein).

### Modeling: Global protein structure modeling

Protein structure modeling was performed using the RoseTTAFold web server as the default modeling program. This is a deep learning-based program that uses a three-track neural network for accurate, rapid structure prediction (Baek et al. 2021) (https://robetta.bakerlab.org/) (**Figure 1A**). Amino acid sequences of all 990 *T. pallidum* proteins were downloaded in FASTA format from NCBI (https://www.ncbi.nlm.nih.gov/nuccore/521094209?sat=56&satkey=64496744) and submitted to RoseTTAFold. Protein structure models were generated for each *T. pallidum* protein sequence submitted to the server, together with a predicted Global Distance Test (GDT) confidence score for each protein. RoseTTAFold GDT scores range from 0.0-1.0, where a greater score is indicative of a more accurate protein structure model. The coordinate file of each RoseTTAFold structure model was downloaded in PDB format with “all coordinates” selected (includes modeled amino acid residues with greater than 5 Å error estimates). Proteins that could not be modeled in RoseTTAFold due to low-confidence, protein size constraints (protein sizes outside the accepted amino acid size range of 26-1200 residues), or data processing errors, were downloaded as PDB files from the AlphaFold database (https://alphafold.ebi.ac.uk/) (Varadi et al. 2022; Varadi et al. 2024). If AlphaFold database models were not available, structures were predicted using the template-based modeling (TBM) web server, I-TASSER (Iterative Threading ASSEmbly Refinement) (https://zhanggroup.org/I-TASSER/) (Yang et al. 2015; Yang and Zhang 2015a; 2015b). Of the top five models predicted for each protein by I-TASSER, the PDB coordinate file corresponding to the highest confidence model was downloaded for each *T. pallidum* protein. I-TASSER model confidence was assessed using the TM-score, a length-independent normalized score ranging from 0 to 1.0, where scores greater than 0.5 indicate correct overall topology. All RoseTTAFold, AlphaFold, and I-TASSER structure modeling in the present study was performed between the years 2022-2025. Protein structure model visualization, labelling, and comparative structure analyses were performed using the molecular visualization programs PyMOL 3.2.0a (https://pymol.org/) (Schrodinger 2015) and UCSF ChimeraX 1.5 (https://www.cgl.ucsf.edu/chimerax/) (Meng et al. 2023).

### Comparative structure analyses: DALI analyses and identification of high-confidence *T. pallidum* protein structure model orthologs

To identify proteins with potential tertiary structural similarity to *T. pallidum* proteins, the PDB coordinate files of the *T. pallidum* structure models generated by RoseTTAFold, the AlphaFold database, and I-TASSER, were submitted to the DALI protein structure comparison server (http://ekhidna2.biocenter.helsinki.fi/dali/) (Holm 2020a; 2020b; 2022; Holm and Sander 1995) (**Figure 1A**) between the years 2022-2025. Due to the DALI alignment score being a continuous scale of structural similarity, length-dependent rescaling of the DALI-score is used to generate a Z-score, with a higher Z-score signifying higher structural similarity (Holm 2020a; 2020b).

Heuristic PDB (Berman et al. 2000) searches were performed in DALI for each *T. pallidum* protein structure model, where the *T. pallidum* query structure was compared against all solved protein structures in the PDB. The output data from the DALI “Full PDB” searches produced lists of all matched proteins with solved structures in the PDB that exhibit varying levels of structural similarity with the *T. pallidum* query structure, in which structural orthologs were ranked from most similar (highest DALI Z-score) to least similar (lowest DALI Z-score). The lowest Z-score outputs in each DALI PDB search were defaulted to 2.0; Z-scores equal to or greater than 2.0 denote significant similarities, and indicate that the query and matched structural ortholog protein from DALI adopt similar folds (Holm et al. 2008). To identify high-confidence *T. pallidum* protein structure model orthologs, the raw, unedited DALI PDB search results were filtered using established minimum confidence thresholds, if available. Structural ortholog matches were considered high-confidence if they met all of the following criteria: RoseTTAFold GDT score (“RoseTTAFold confidence”) ≥0.50 (equivalent to 50% confidence), reflecting the overall modeling confidence across entire proteins, including unstructured regions and extracellular loops that are difficult to model; DALI Z-score (“Z-Score”) ≥ 8.0, ensuring a high probability of structural homology (Holm 2020b); alignment coverages (the number of structurally equivalent aligned residue pairs) of the query protein (“*T. pallidum* Protein Query Coverage”) and the PDB structural ortholog (“Protein Template Coverage”) both ≥10%, ensuring the inclusion of small functional domains that have been solved and deposited in the PDB (Houston et al. 2018); and structural resolution of the PDB ortholog (“PDB Resolution”) <4.0Å, which excluded protein structures where only some chains are solved and when domain folds and large structural features are the highest resolved structural features. This resolution criteria cutoff was based on the guide for interpreting values of resolution in FirstGlance in Jmol version 16.1, an open-source macromolecular visualization software package (http://firstglance.jmol.org). The alignment coverages of the query and PDB structural ortholog proteins were calculated using the following formulae:

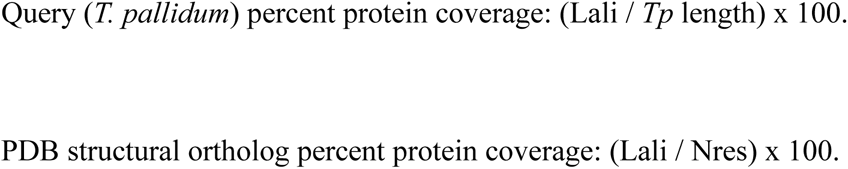

Lali: the number of structurally equivalent aligned residue pairs (Holm 2020a; 2020b). Nres: the total number of amino acids in the PDB structural ortholog (Holm 2020a; 2020b).

### Protein annotation: Structure homology-based assignment of potential protein functions

#### (1) Proteome annotation confirmation

In order to validate our structure modeling-to-function method, the level of correlation between *T. pallidum* NCBI proteome annotations (functionally annotated proteins only) and our corresponding structure modeling-based annotations was determined (**Figure 1B**). In the present study, the predicted functions of *T. pallidum* proteins were assigned based on the protein functions (PDB molecule description outputs in DALI) of the high-confidence structural ortholog matches (higher than the minimum threshold cutoff values listed above) from the DALI PDB searches. These putative structure modeling-based protein functions were then compared with the genome sequence-based functional annotations of the *T. pallidum* proteins in the NCBI proteome. For calculating the level of correlation between the sequence- and structure-based annotation methods, all functionally annotated *T. pallidum* proteins were categorized as either (1) “same function”, (2) “related function”, or (3) “different function”.

#### (2) Multi- and novel domain analyses

A strategy was developed to address the complexity of multidomain architectures in *T. pallidum* proteins by identifying multidomain proteins with unique or potentially novel domain combinations, in order to improve the functional annotation of full-length proteins that may contain multiple domains and functions. Full-length amino acid sequences were first divided into individual domain sequences using DALI pairwise amino acid sequence alignments to define domain boundaries. Each domain sequence was then analyzed separately using the structure modeling-to-function workflow. To identify functionally annotated *T. pallidum* multi-domain proteins that contain unique domain combinations, proteins were filtered to retain only those proteins whose predicted domains have different functions (**Figure 1C**). Literature reviews and PDB searches were then performed to identify domain combinations that, to our knowledge, have not been previously described in the literature or deposited in the PDB. To identify functionally annotated *T. pallidum* proteins with potentially novel domains, DALI PDB search results were examined for domains that were successfully modeled by RoseTTAFold but lacked structurally orthologous domains (no DALI matches or Z-scores below 2.0).

#### (3) Novel protein functional annotation

A search of the *T. pallidum* Nichols proteome identified 323 proteins of unknown function. These included 149 “hypothetical proteins”, 32 DUF proteins, and 142 proteins with annotations that provide minimal insight into function. We analyzed the functions of the high-confidence structural ortholog matches that were identified for each of these 323 treponemal proteins (**Figure 1C**). This approach used the same DALI PDB searches and minimum threshold cutoff values for the identification of high-confidence structural orthologs as outlined above.

### Functional categorization of *T. pallidum* proteins

The genome-wide functional annotation tool, eggNOG-mapper (version 2.1.9) (http://eggnog-mapper.embl.de/) (Cantalapiedra et al. 2021), was used to assign *T. pallidum* proteins to functional categories based on the “Clusters of Orthologous Genes” (COG) (Tatusov et al. 1997) categories (https://www.ncbi.nlm.nih.gov/research/cog). Searches were performed using the eggNOG 5 database (Huerta-Cepas et al. 2019), default search parameters, and default annotation options.

### OMP analyses and B cell epitope mapping

Given that OMPs exposed on the host-facing surface of *T. pallidum* are likely determinants of virulence and represent key targets for syphilis vaccine design (Hawley et al. 2021; Izard et al. 2009; Liu et al. 2024; Liu et al. 2010; Radolf and Kumar 2017), we undertook a systematic computational analysis of known and putative *T. pallidum* OMPs to model and characterize their structures using multiple AI-based modeling programs, assess the confidence of the OMP designation, and identify and prioritize predicted surface-exposed BCEs. These analyses were comprised of three main components (**Figure 1C**):

### (1) Additional protein structure modeling

To obtain high-confidence structure models of known and putative *T. pallidum* OMPs for downstream analyses, additional structure predictions were performed using AlphaFold 3 as follows. Mature amino acid sequences of the known/predicted *T. pallidum* OMPs were generated by the removal of N-terminal signal peptides predicted by the signal peptide prediction server, SignalP 6.0 (https://services.healthtech.dtu.dk/services/SignalP-6.0/) (Teufel et al. 2022). Mature OMP sequences were then submitted to the AlphaFold 3 server (https://alphafoldserver.com/) (Abramson et al. 2024). AlphaFold 3 protein structure models were generated for each *T. pallidum* protein sequence, together with confidence scores for each model: (1) pLDDT (predicted local distance difference test) score which gives a per-atom confidence estimate on a 0-100 scale where a higher value indicates higher confidence; (2) PAE (predicted aligned error) which estimates the error in the relative position and orientation of two amino acid residues in the predicted structure, and; (3) pTM (the predicted template modeling) and ipTM (interface predicted template modeling) scores, which are measures of whole molecule structural accuracy. pLDDT scores were normalized to a [0,1] scale for consistency with RoseTTAFold and I-TASSER. pLDDT and pTM scores of 0.5 or greater indicate higher quality predictions. ipTM scores greater than 0.8 indicate high-confidence predictions, whereas scores lower than 0.6 indicate unreliable structure predictions. The coordinate files for the highest confidence AlphaFold 3 models were used for visualization, labelling, comparative analyses, and all downstream bioinformatics and epitope analyses in the molecular visualization systems, PyMOL 3.2.0a (https://pymol.org/) (Schrodinger 2015) and UCSF ChimeraX 1.5 (https://www.cgl.ucsf.edu/chimerax/) (Meng et al. 2023).

### (2) Structure- and sequence-based bioinformatics analyses

Bioinformatic characterization of known and putative *T. pallidum* OMPs was performed using complementary structure- and sequence-based analyses to assess the likelihood that these proteins are true OMPs and to map their host-facing surfaces for downstream epitope mapping. (a) The presence of cleavable signal peptide sequences was predicted using the following web servers: SignalP 6.0 (https://services.healthtech.dtu.dk/services/SignalP-6.0/) (Teufel et al. 2022), DeepSig (http://busca.biocomp.unibo.it/deepsig/) (Savojardo et al. 2018), PrediSi (http://www.predisi.de/) (Hiller et al. 2004), Phobius (https://phobius.sbc.su.se/) (Kall et al. 2007), and TOPCONS 2.0 (including TOPCONS, Philius, PolyPhobius, and SPOCTOPUS prediction methods) (https://topcons.net/) (Tsirigos et al. 2015). For greater stringency, two additional web server algorithms were used to predict signal peptides in the putative novel OMPs identified in the present study: Signal-CF (http://www.csbio.sjtu.edu.cn/bioinf/Signal-CF/) (Chou and Shen 2007) and Signal-3L 3.0 (http://www.csbio.sjtu.edu.cn/bioinf/Signal-3L/) (Zhang et al. 2020). (b) Protein subcellular localization was predicted using the following web servers: DeepLocPro 1.0 (https://services.healthtech.dtu.dk/services/DeepLocPro-1.0/) (Moreno et al. 2024), PSORTb version 3.0.3 (https://www.psort.org/psortb/) (Yu et al. 2010), and CELLO 2.5 (http://cello.life.nctu.edu.tw/) (Yu et al. 2006). (c) Beta-barrel OMPs were predicted using DeepTMHMM 1.0.42 (https://dtu.biolib.com/DeepTMHMM) (J. Hallgren 2022) and PRED-TMBB (http://bioinformatics.biol.uoa.gr/PRED-TMBB/) (Bagos et al. 2004) web servers. (d) Mapping predicted surface-exposed, host-facing regions of known and potential treponemal OMPs was performed using DeepTMHMM to determine the membrane topology of these beta-barrel domain-containing OMPs. Amino acids predicted by DeepTMHMM to be located in “outside” (extracellular environment), “membrane” (outer membrane-embedded), or “inside” (periplasm) subcellular locations were used to determine the overall orientation of each protein in the outer membrane. Known and potential OMP structure models were orientated with “inside” and “outside” regions positioned at the bottom and top of the protein structure models, respectively. Putative surface-exposed, host-facing protein regions (including extracellular loops, ECLs) were then mapped via the identification of all amino acid sequences in predicted “outside” regions.

### (3) Putative B cell epitope identification, mapping, and prioritization

To identify vaccine-relevant targets, we next sought to identify and prioritize BCEs in known and putative *T. pallidum* OMPs. Four prediction programs were used for the identification of putative linear BCEs in known and potential OMPs from *T. pallidum*: (i) ElliPro (protein structure-based method focused on geometric properties: http://tools.iedb.org/ellipro/) (Ponomarenko et al. 2008); (ii) ABCPred (machine learning discrimination model using Artificial Neural Networks [ANNs]: https://webs.iiitd.edu.in/raghava/abcpred/index.html) (Saha and Raghava 2006); (iii) BepiPred 3.0 (machine learning discrimination model using ANNs: https://services.healthtech.dtu.dk/services/BepiPred-3.0/) (Clifford et al. 2022); and (iv) SVMTriP (machine learning discrimination model using Support Vector Machine [SVM]: http://sysbio.unl.edu/SVMTriP/) (Yao et al. 2012). Coordinate files of the *T. pallidum* protein structure models based on the SignalP-prediction results were submitted to ElliPro, and the corresponding amino acid sequences were submitted to ABCPred, BepiPred, and SVMTriP, all using default settings. These analyses were all performed between November 2024 and April 2025. The location of each amino acid sequence corresponding to the predicted linear BCEs from each of the four prediction servers were mapped onto the RoseTTAFold and AlphaFold 3 structure models of each known/putative *T. pallidum* OMP using the PyMOL molecular visualization tool. The first step in prioritizing and ranking predicted BCEs was determining whether the epitopes are located in surface-exposed regions of the known and potential OMP structure models, or in non-surface-exposed regions (sites buried within the outer membrane or oriented towards the periplasm). Only potential epitopes that contain at least five amino acids located within predicted surface-exposed, host-facing regions were analyzed further (Kringelum et al. 2013; Singh et al. 2013). High-ranking predicted BCEs were identified based on the following criteria: (1) predicted by at least two prediction servers; (2) prediction scores of 50% or higher; (3) lacked cysteine residues; (4) predominantly composed of hydrophilic residues (negative or near-neutral Grand Average of Hydropathy [GRAVY] scores; -3.0 to +0.5, highly hydrophilic to mildly hydrophobic); (5) conserved across all NCBI *T. pallidum* strains; (6) exhibited ≤ 75% sequence identity with human proteins; (7) protein ranked as a high-/medium-confidence OMP, as described above, and; (8) protein expression detected in previous *T. pallidum* studies (Houston et al. 2023; Houston et al. 2024a; Osbak et al. 2016). Amino acid conservation analyses were performed using NCBI PSI-BLAST searches (default settings) against *T. pallidum* ssp. *pallidum* proteins (taxid:161) and human proteins (taxid:9606) (https://blast.ncbi.nlm.nih.gov/Blast.cgi?CMD=Web&PAGE=Proteins&PROGRAM=blastp&RUN_PSIBLAST=on) (Altschul et al. 1997).

### Data integration: comparative analyses of SS14-Nichols inter-strain protein sequence differences

Protein sequence alignments of seven potential novel OMPs from Nichols and SS14 (NC_021508) strains were conducted using EMBOSS Needle (https://www.ebi.ac.uk/jdispatcher/psa/emboss_needle) (Madeira et al. 2024) to identify amino acid differences between orthologous proteins. Mass spectrometry (MS)-based proteomics data from previous *T. pallidum* studies (Houston et al. 2023; Houston et al. 2024a; Houston et al. 2024b; Osbak et al. 2016; Romeis et al. 2021) were analyzed to validate the sequence differences in expressed proteins by identifying tryptic peptides containing the variant residues. RoseTTAFold and AlphaFold 3 were used for structure modeling and mapping of the potential novel OMPs with annotated and experimentally validated inter-strain amino acid differences.

## Acknowledgements

This research was supported by R37AI051334, U19AI144133 and U01AI182035 (CEC) from the National Institute of Allergy and Infectious Diseases (NIAID) at the National Institutes of Health (NIH), 235655 from Coefficient Giving (CEC), as well as 196519 from the Canadian Institutes of Health Research (CIHR) (JAA).

## Data availability statement

Supplementary Figures S1-S9 and Supplementary Tables S1-S17, protein structure modeling coordinate files, structure models, and raw data have been deposited to the Harvard Dataverse repository (https://dataverse.harvard.edu/) and are available using the following link: https://dataverse.harvard.edu/dataset.xhtml?persistentId=doi:10.7910/DVN/85TKGN.

## Author Contributions

**Simon Houston:** Conceptualization, writing – original draft, methodology, investigation, formal analyses, data curation, visualization, writing – review and editing, supervision. **Steven Marshall:** Methodology, investigation, formal analyses, data curation, visualization, writing - review and editing. **Austin Miller:** Methodology, investigation, formal analysis, visualization. **Aleksander Palkowski:** Investigation, formal analyses, writing - review and editing. **Javier A. Alfaro:** Investigation, formal analyses, writing - review and editing. **Caroline E. Cameron:** Conceptualization, formal analyses, writing - review and editing, funding acquisition, project administration, supervision.

